# Generalised free energy and active inference: can the future cause the past?

**DOI:** 10.1101/304782

**Authors:** Thomas Parr, Karl J Friston

## Abstract

We compare two free energy functionals for active inference under Markov decision processes. One of these is a functional of beliefs about states and policies, but a function of observations, while the second is a functional of beliefs about all three. In the former (*expected* free energy), prior beliefs about outcomes are not part of the generative model (because they are absorbed into the prior over policies). Conversely, in the second (*generalised* free energy); priors over outcomes become an explicit component of the generative model. When using the free energy function, which is blind to counterfactual (i.e., future) observations, we equip the generative model with a prior over policies that ensure preferred (i.e., priors over) outcomes are realised. In other words, selected policies minimise uncertainty about future outcomes by minimising the free energy expected in the future. When using the free energy functional – that effectively treats counterfactual observations as hidden states – we show that policies are inferred or selected that realise prior preferences by minimising the free energy of future expectations. Interestingly, the form of posterior beliefs about policies (and associated belief updating) turns out to be identical under both formulations, but the quantities used to compute them are not.

## 1. Introduction

Over the past years, we have tried to establish active inference (a corollary of the free energy principle) as a relatively straightforward and principled explanation for action, perception and cognition. Active inference can be summarised as self-evidencing (Hohwy 2016); in the sense that action and perception can be cast as maximising Bayesian model evidence, under generative models of the world. When this maximisation uses approximate Bayesian inference, this is equivalent to minimising variational free energy (Friston et al. 2006) – a form of bounded rational behaviour that minimises a variational bound on model evidence. Recently, we have migrated the basic idea from models that generate continuous sensations (like velocity and luminance contrast) (Brown and Friston 2012) to discrete state-space models; specifically Markov decision processes (Friston et al. 2017a). These models represent the world in terms of discrete states; like I am on this page and reading this word (Friston et al. 2017c). Discrete state-space models can be inferred using belief propagation (Yedidia et al. 2005) or variational message passing (Dauwels 2007; Winn 2004) schemes that have a degree of neuronal plausibility (Friston et al. 2017b). The resulting *planning as inference* scheme (Attias 2003; Baker et al. 2009; Botvinick and Toussaint 2012; Verma and Rao 2006) has a pleasingly broad explanatory scope; accounting for a range of phenomena in cognitive neuroscience, active vision and motor control (see Table 1). In this paper, we revisit the role of (expected) free energy in active inference and offer an alternative, simpler and more general formulation. This formulation does not substantially change the message passing or belief updating; however, it provides an interesting perspective on planning as inference and the way that we may perceive the future.

**Table 1:** applications of active inference for Markov decision processes

| Application | Comment | References |
| --- | --- | --- |
| **Decision making under uncertainty** | Initial formulation of active inference for *Markov decision processes* and *sequential policy optimisation* | (Friston et al. 2012c) |
| **Optimal control (the mountain car problem)** | Illustration of *risk sensitive or KL control* in an engineering benchmark | (Friston et al. 2012a) |
| **Evidence accumulation: Urns task** | Demonstration of how beliefs states are absorbed into a generative model | (FitzGerald et al. 2015b; FitzGerald et al. 2015c) |
| **Addiction** | Application to psychopathology | (Schwartenbeck et al. 2015c) |
| **Dopaminergic responses** | Associating dopamine with the encoding of (expected) precision provides a plausible account of dopaminergic discharges | (FitzGerald et al. 2015a; Friston et al. 2014) |
| **Computational fMRI** | Using Bayes optimal precision to predict activity in dopaminergic areas | (Schwartenbeck et al. 2015a) |
| **Choice preferences and epistemics** | Empirical testing of the hypothesis that people prefer to keep options open | (Schwartenbeck et al. 2015b) |
| **Behavioural economics and trust games** | Examining the effects of prior beliefs about self and others | (Moutoussis et al. 2014) |
| **Foraging and two step mazes; navigation in deep mazes** | Formulation of epistemic and pragmatic value in terms of *expected free energy* | (Friston et al. 2015) |
| **Habit learning, reversal learning and devaluation** | Learning as minimising variational free energy with respect to model parameters – and action selection as *Bayesian model averaging* | (FitzGerald et al. 2014; Friston et al. 2016) |
| **Saccadic searches and scene construction** | *Mean field approximation* for multifactorial hidden states, enabling high dimensional beliefs and outcomes: c.f., functional segregation | (Friston and Buzsaki 2016; Mirza et al. 2016) |
| **Electrophysiological responses: place-cell activity, omission related responses, mismatch negativity, P300, phase-precession, theta-gamma coupling** | Simulating neuronal processing with a gradient descent on variational free energy; c.f., dynamic *Bayesian belief propagation* based on marginal free energy | (Friston et al. 2017a) |
| **Structure learning, sleep and insight** | Inclusion of parameters into expected free energy to enable structure learning via *Bayesian model reduction* | In press |
| **Narrative construction and reading** | Hierarchical generalisation of generative model with *deep temporal structure* | (Friston et al. 2017c) |

In current descriptions of active inference, the basic argument goes as follows: active inference is based upon the maximisation of model evidence or minimisation of variational free energy in two complementary ways. First, one can update one’s beliefs about latent or hidden states of the world to make them consistent with observed evidence – or one can actively sample the world to make observations consistent with beliefs about states of the world. The important thing here is that both action and perception are in game of minimising the same quantity; namely, variational free energy. A key aspect of this formulation is that action (i.e., behaviour) is absorbed into inference, which means that agents have beliefs about what they are doing – and will do. This calls for prior beliefs about action or policies (i.e., sequences of actions). So where did these prior beliefs come from?

The answer obtains from a *reductio ad absurdum* argument: if action realises prior beliefs and minimises free energy, then the only tenable prior beliefs are that action will minimise free energy. This leads to the prior belief that I will select policies that minimise the free energy expected under that policy. The endpoint of this argument is that action or *policy selection becomes a form of Bayesian model selection*, where the evidence for a particular policy becomes the free energy expected in the future. This *expected free energy* is a slightly unusual objective function because it scores the evidence for plausible policies based on outcomes that have yet to be observed. This means that the expected free energy becomes the variational free energy expected under (posterior predictive) beliefs about outcomes. These priors are usually informed by prior beliefs about outcomes that play the role of prior preferences or utility functions in reinforcement learning and economics.

In summary, beliefs about states of the world and policies are continuously updated to minimise variational free energy, where posterior beliefs about policies (that prescribe action) are based upon expected free energy (that may or may not include prior preferences over future outcomes). This is the current story and leads to interesting issues that rest on the fact that expected free energy can be decomposed into epistemic and pragmatic parts (Friston et al. 2015). This decomposition provides a principled explanation for the epistemics of planning and inference that underwrite the exploitation and exploration dilemma, novelty, salience and so on. However, there is another way of telling this story that leads to a conceptually different sort of interpretation.

In what follows, we show that the same Bayesian policy (model) selection obtains from minimising variational free energy when *future outcomes are treated as hidden or latent states of the world*. In other words, we can regard active inference as minimising a generalised free energy under generative models that entertain the consequences of (policy-dependent) hidden states of the world in the future. This simple generalisation induces posterior beliefs over future outcomes that now play the role of counterfactual, latent or hidden states. In this setting, the future is treated in exactly the same way as the hidden or unobservable states of the world generating observations in the past. On this view, one gets the expected free energy for free, because the variational free energy involves an expectation under posterior beliefs over future outcomes. In turn, this means that beliefs about states and policies can be simply and uniformly treated as minimising the same (generalised) free energy, without having to invoke any free energy minimising priors over policies.

Technically, this leads to the same form of belief updating and (Bayesian) policy selection but provides a different perspective on the free energy principle *per se*. This perspective says that self-evidencing and active inference both have one underlying imperative; namely, to minimise *generalised free energy* or uncertainty. When this uncertainty is evaluated under models that generate outcomes in the future, future outcomes become hidden states that are only revealed by the passage of time. Formally, the ensuing generalised free energy is a Hamiltonian Action, because it is a path or time integral of free energy at each time point. In other words, active inference is just a statement of Hamilton’s Principle of Stationary Action. In this context, outcomes in the past become observations in standard variational inference, while outcomes in the future become posterior beliefs about latent observations that have yet to disclose themselves. In this way, the generalised free energy can be seen as comprising variational free energy contributions from the past and future.

The current paper provides the formal basis for the above arguments. In brief, we will see that both the expected and generalised free energy formulations lead to the same update equations. However, there is a subtle difference. In the expected free energy formalism, prior preferences or beliefs about outcomes are used to specify the prior over policies. In the generalised formulation, prior beliefs about outcomes in the future inform posterior beliefs about the hidden states that cause them. Because of the implicit forward and backward message passing in the belief propagation scheme, these prior beliefs or preferences act to distort expected trajectories (into the future) towards preferences in an optimistic way (Sharot et al. 2012). Intuitively, the expected free energy contribution to generalised free energy evaluates the (complexity) cost of this distortion; thereby favouring policies that lead naturally to preferred outcomes – without violating beliefs about state transitions and the (likelihood) mapping between states and outcomes. The implicit coupling between beliefs about the future and current actions means that, in one sense, the future can cause the past.

This paper comprises three sections. In the first, we outline the approach we have used to date (i.e., minimising the variational free energy under prior beliefs that policies with a low expected free energy are more probable). In the second, we introduce a generalisation of the variational free energy that incorporates beliefs about counterfactual outcomes. The third section compares these two approaches conceptually and through illustrative simulations.

## 2. Active inference and variational free energy

The free energy principle is motivated by the defining characteristic of living creatures; namely, that they persist in the face of a changing world. In other words, their states occupy a small proportion of all possible states with a high probability. Mathematically, this means that they show a form of self organised, non-equilibrium steady-state that maintains a low entropy probability distribution over their states. In information theory, self information or *surprise* (a.k.a. negative log model evidence) averaged over time is entropy. This means, at any given time, all biological systems are compelled to minimise their surprise. Although the computation of surprise is often intractable, an approximation is simple to calculate. This is variational free energy (Beal 2003; Dayan et al. 1995; Friston 2003) which, as Jensen’s inequality demonstrates, is an upper bound on surprise.

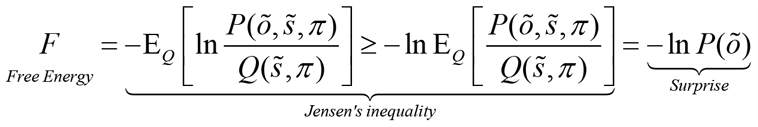

In the equation above, *P* indicates a probability distribution over outcomes that are generated by hidden states of the world, which defines the system’s generative model. *Q* is a probability distribution over unobservable (hidden) states that becomes an approximate posterior distribution as free energy is minimised. The minimisation of free energy over time ensures entropy does not increase, thereby enabling biological systems to resist the second law of thermodynamics and their implicit dissipation or decay. Active inference is the process of reducing free energy through action and perception.

In the following, we begin by describing the form of the generative model we have used to date. We will then address the form of the approximate posterior distribution. To make inference tractable, this reform generally involves a mean-field approximation that factorises the approximate posterior distribution into independent factors or marginal distributions.

The generative models used in this paper are subtly different for each free energy functional, but the variables themselves are the same. These are policies, ***π***, and states at different times: 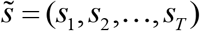, all of which are latent (unknown random) variables that have to be inferred. States evolve as a discrete Markov chain, where the transition probabilities are functions of the policy. Likelihood distributions probabilistically map hidden states to observations: 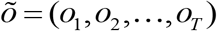. Figure 1 (left) shows these dependencies as a graphical Bayesian network. This type of generative model has been used extensively in simulations of active inference (FitzGerald et al. 2014; FitzGerald et al. 2015c; Friston et al. 2017a; Friston et al. 2015; Friston et al. 2017b; Friston et al. 2017c; Schwartenbeck et al. 2015a): please see Table 1.

**Figure 1 -.**
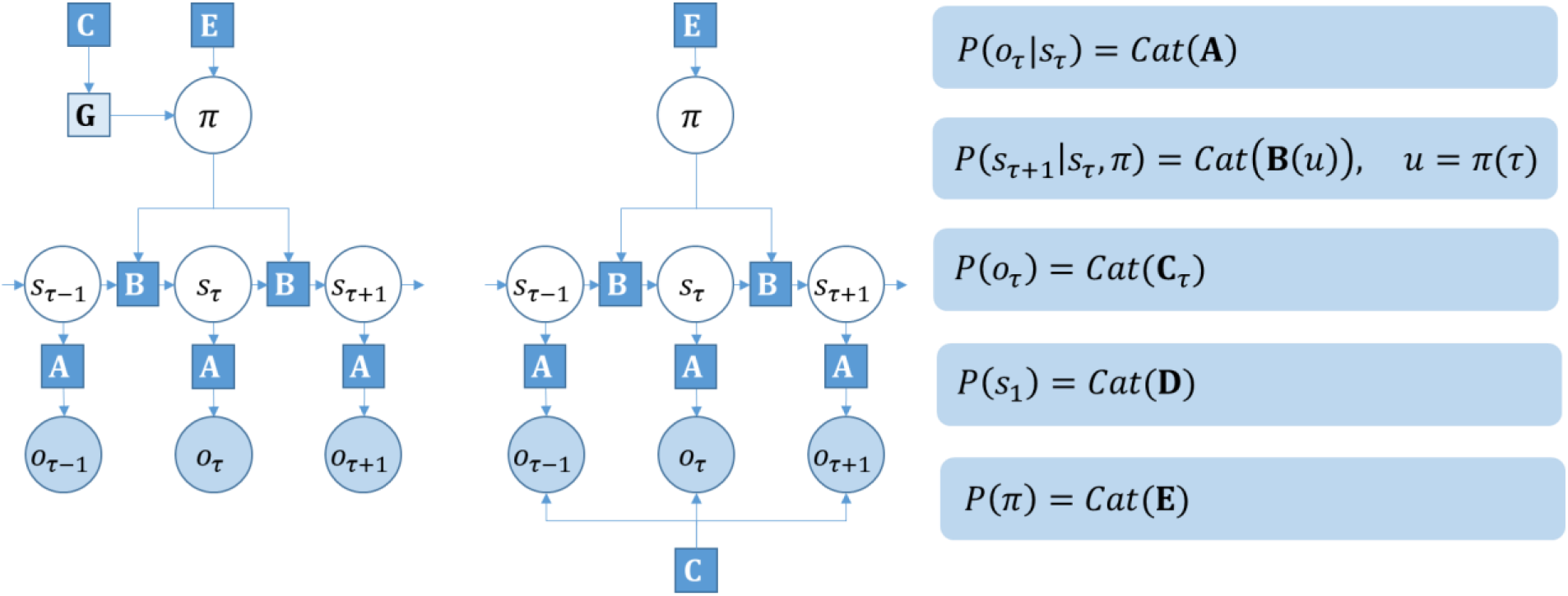
Markov decision process. This shows the basic structure of the discrete state space generative model used in this paper. The factor graph *on the left* is the generative model we have used in previous work. Importantly, the prior belief about observations only enters this graph through the expected free energy, *G* (see main text), which enters the prior over policies. The *right* factor graph is the new version of the generative model considered in this paper. This generative model does not require an expected free energy, and the prior over outcomes enters the model directly as a constraint on outcomes. Please refer to the main text and Table 2 for a description of the variables. In the panels *on the right*, the definitions are given for each of the factors in blue squares. Here, Cat refers to the categorical distribution.

**Table 2:**
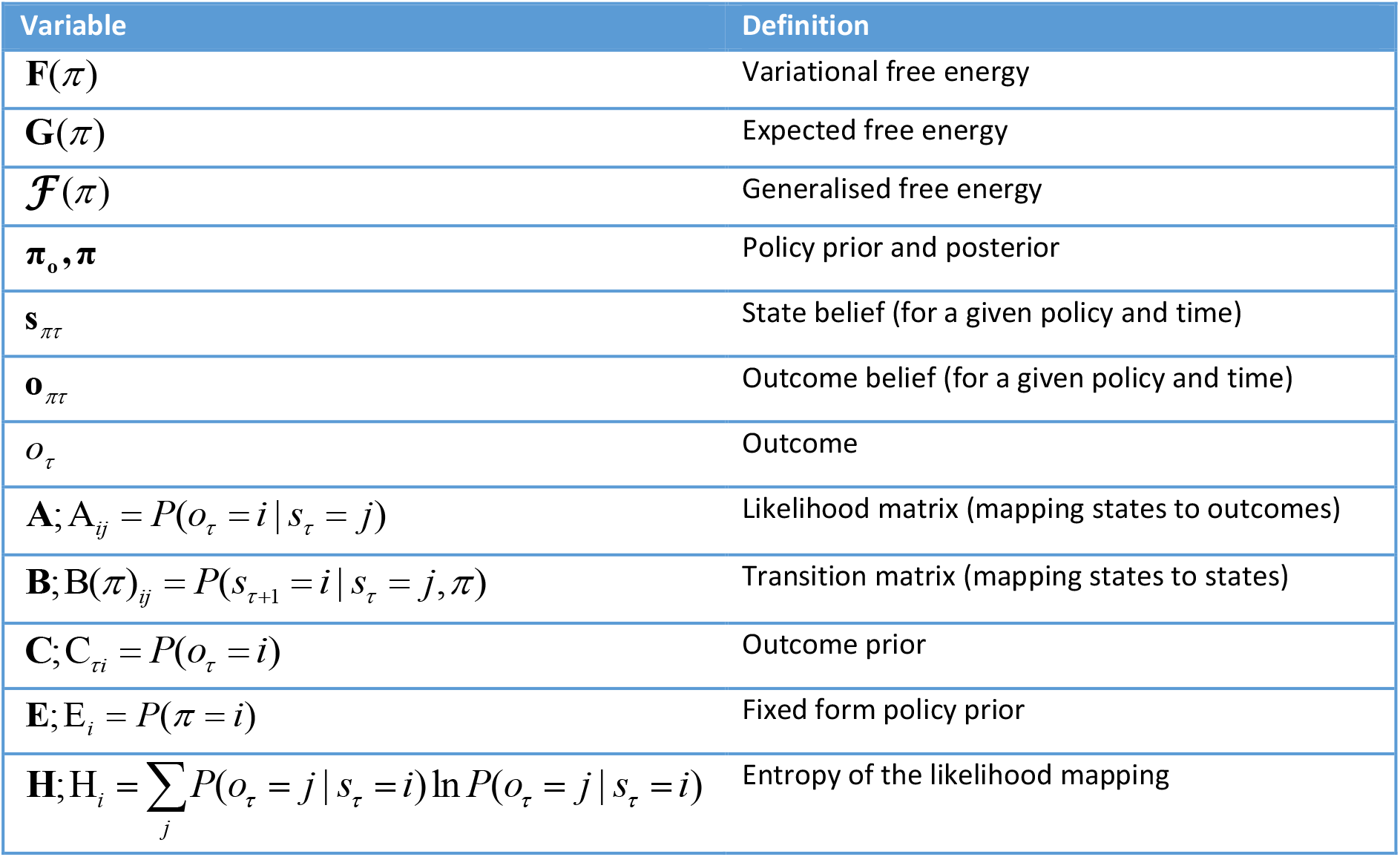
Variables in update equations

It is worth noting that the free energy is a *functional* of the distributions in the generative model, and of the approximate posterior beliefs, but a *function* of observations. Continuing with this free energy, we now consider the mean field approximation in current implementations of active inference, and its consequences for the variational free energy.

### 2.1 Definition of the variational free energy

To define the variational free energy for the above generative model, we first need to specify the form of the approximate posterior distribution, *Q*. We do this via a mean field approximation that treats the (policy dependent) state at each time step as approximately independent of the state at any other time step. We treat the distribution over the policy as a separate factor, which implies a set of models, ***π***, over hidden variables *s_τ_*:

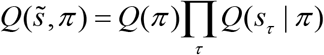

Substituting this in to the free energy definition above, we get the variational free energy:

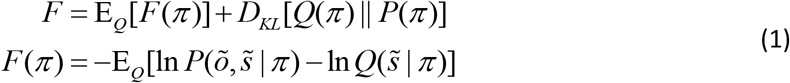

In this form, the variational free energy is expressed in terms of policy dependent terms (second equality) that bound the (negative log) evidence for each policy and a complexity cost or KL divergence that scores the departure of the posterior beliefs over policies from the corresponding prior beliefs.

### 2.2 Past and Future

There is an important difference in how past and future outcomes are treated by the variational free energy. Note that – as a function of outcomes – the components of the free energy that depend on outcomes can only be evaluated for the past and present. Hidden states, on the other hand, enter the expression as *beliefs* about states. In other words, the free energy is a functional of distributions over states, rather than a function, as in the case of outcomes. This means that free energy evaluation takes account of future states. We can express this explicitly by writing the variational free energy as a sum over time, factorising the generative distribution according to the conditional independencies expressed in Figure 1 (left):

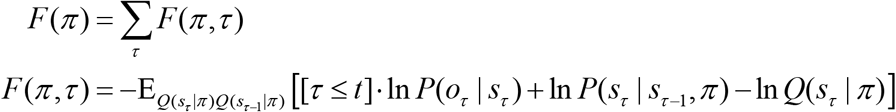

In the above, the Iverson (square) brackets return 1 if the expression is true, and 0 otherwise. It is this condition that differentiates contributions from the past from the future. This allows us to decompose the sum into past and future components:

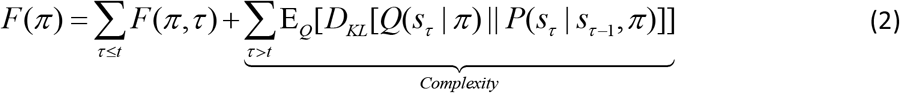

In this decomposition, the contribution of beliefs about future states reduces to a complexity cost that scores the KL divergence between approximate posterior beliefs about states in the future, relative to the prior beliefs based upon the (policy-specific) transition probabilities in the generative model.

### 2.3 Policy posteriors and pSh2 riors

Using the full variational free energy (over all policies) from Equation 1, we can evaluate posterior beliefs about policies. The variational derivative of the free energy with respect to these beliefs is (where σ(·) is a softmax function):

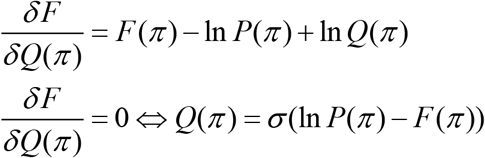

This, together with Equation 2, implies the belief prior to any observations (i.e., at *τ* = 0), which is given by:

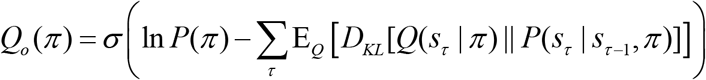

This is an unsatisfying result, in that it fails to accommodate our prior knowledge that outcomes will become available in the future. In other words, the posterior at each time step is calculated under a different model (see Figure 2).

**Figure 2 -.**
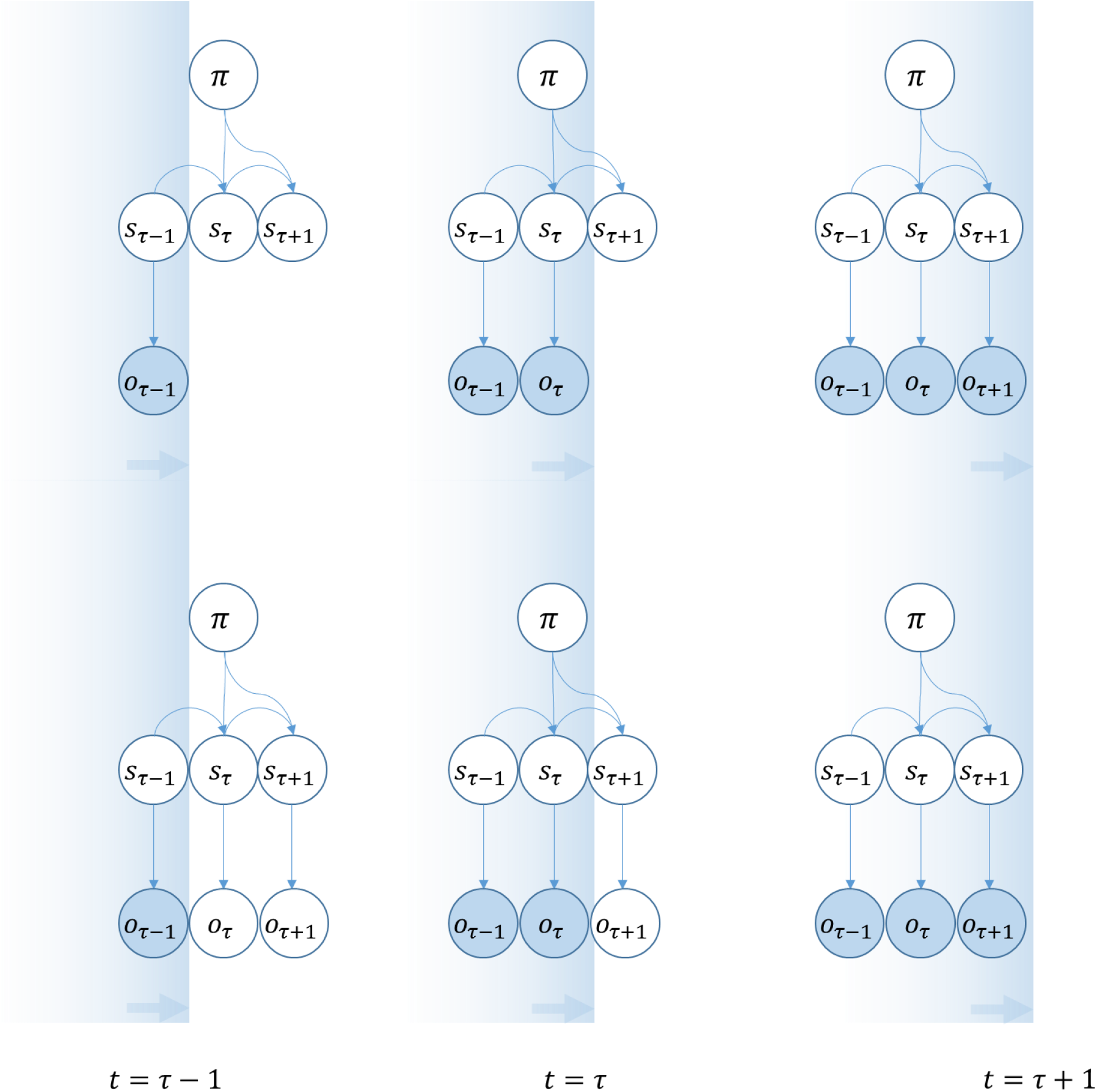
Temporal progression of MDP. The upper graphs shows the structure of the generative model implied using the variational free energy, equipped with a prior that the expected free energy will be minimised by policy selection. Observations are added to the model as they occur. The lower graphs show the structure of the generative model that explicitly represents counterfactual outcomes, and minimises a generalised free energy through policy selection. As observations are made, the outcome variables collapse to delta functions.

### 2.4 Expected free energy

To finesse this shortcoming we can assume agents select the policy that they expect will lead to the lowest free energy (summed over time). This is motivated by the *reductio ad absurdum* in the introduction, and is expressed mathematically as:

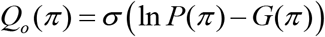

(We retain the notation *Q_o_* (*π*) for the prior here to distinguish this from the fixed form prior *P*(*π*), which does not depend on the beliefs about states). *G*(*π*) is the expected free energy, conditioned on a policy. It is defined as:

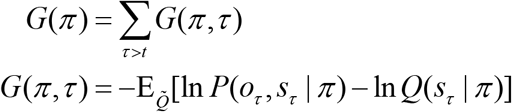

There is an apparent problem with this quantity: the first term within the expectation is a function of outcomes that have yet to be observed. To take this into account, we have defined an (approximate) joint distribution over states and outcomes: 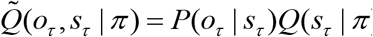, and take the expectation with respect to this. This means that we can express a (posterior predictive) belief about the observations in the future based on (posterior predictive) beliefs about hidden states. One can obtain a useful form of the expected free energy by rearranging the above: if we factorise the generative model, we obtain:

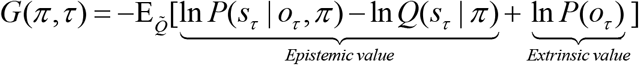

This form shows that policies that have a low expected free energy are those that resolve uncertainty, and that fulfil prior beliefs about outcomes. It is the first of these terms that endorses the metaphor of the brain as a scientist, performing experiments to verify or refute hypotheses about the world (Friston et al. 2012b; Gregory 1980). The second term speaks to the notion of a ‘crooked scientist’ (Bruineberg et al. 2016), who designs experiments to confirm prior beliefs; i.e., preferred outcomes. Through Bayes’ rule,

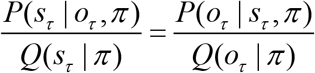

and noting that 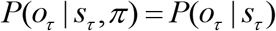, we can also expresses expected free energy in terms of risk and ambiguity:

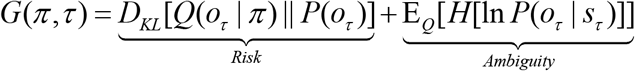

This means that the prior belief about outcomes enters the generative model through the KL-Divergence between outcomes expected under any policy and prior preferences. This form also illustrates the correspondence between the expected free energy and the quantities ‘risk’ and ‘ambiguity’ from behavioural economics (Ellsberg 1961; Ghirardato and Marinacci 2002). Risk quantifies the expected cost of a policy as a divergence from preferred outcomes and is sometimes referred to as Bayesian risk or regret (Huggins and Tenenbaum 2015); which underlies KL control and related Bayesian control rules (Kappen et al. 2012; Ortega and Braun 2010; Todorov 2008) and special cases that include Thompson sampling (Lloyd and Leslie 2013; Strens 2000). Ambiguous states are those that have an uncertain mapping to observations. The greater these quantities, the less likely it is that the associated policy will be chosen.

### 2.5 Hidden state upSh2 dates

To complete our description of active inference, we derive the belief update equations for the hidden states:

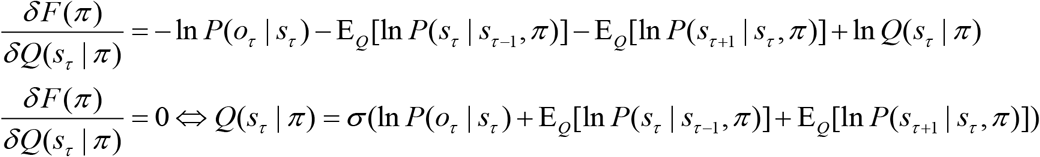

### 2.6 Summary

In the above, we have provided an overview of our approach to date. This uses a variational free energy functional to derive belief updates, while policy selection is performed based on an expected free energy. The resulting update equations are shown in Figure 3 (blue panels). This formulation has been very successful in explaining a range of cognitive functions, as summarised in Table 1. In the following, we present an alternative line of reasoning. As indicated in Figure 2, there is more than one way to think about the data assimilation and evidence accumulation implicit in this formulation. So far, we have considered the addition of new observations as time progresses. We now consider thecase in which (counterfactual) outcomes are represented throughout time. This means that future or latent outcomes have the potential to influence beliefs about past states.

**Figure 3 -.**
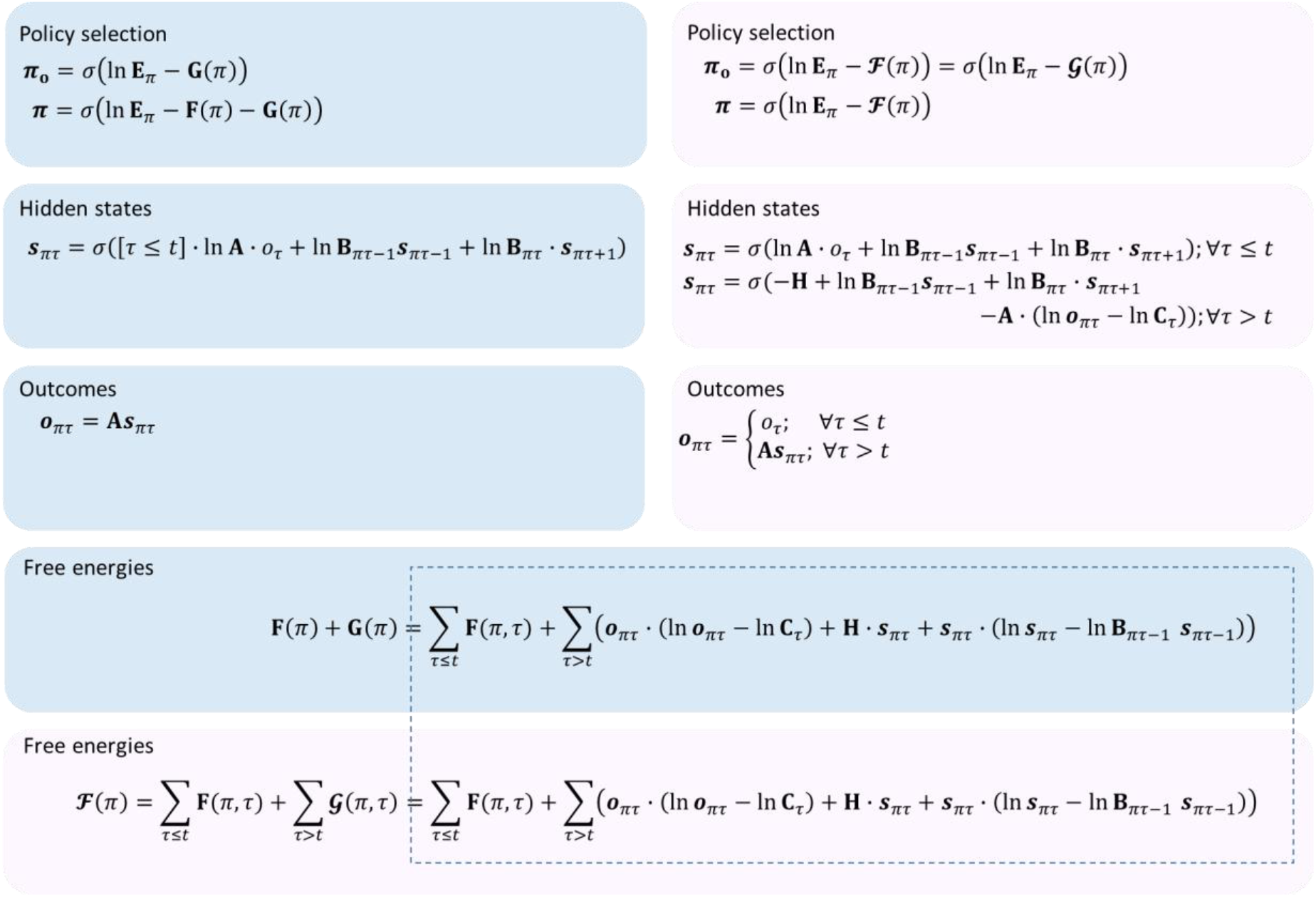
Belief update equations. The blue panels show the update equations using the standard variational approach. The pink panels show the update equations when the generalised free energy is used. The dotted outline indicates the correspondence between the generalised free energy and the sum of the variational and expected free energies, and therefore the equivalence of *the form* of the posteriors over policies. However, it should be remembered that the variables within these equations are not identical, as the update equations demonstrate. See Table 2 for the definitions of the variables as they appear here. The equations used here are discrete updates. A more biologically plausible (gradient ascent) scheme is used in the simulations. These simply replace the updates with differential equations that have stationary points corresponding to the variational solutions above.

## 3. Active inference and generalised free energy

We define the generalised free energy as

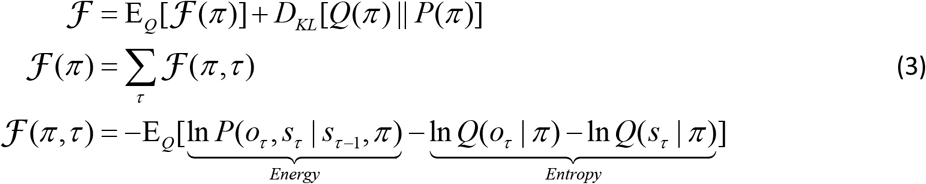

Where, as above, the expectation is with respect to 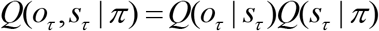. However, we now distinguish the past and the future through the following:

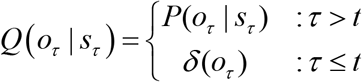

In the generalised free energy, the marginals of the joint distribution over outcomes and states define the entropy but the expectation is over the joint distribution. It is important to note that 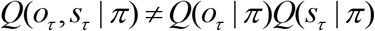. It is this inequality that underlies the epistemic components of generalised free energy. Interestingly, if we assumed conditional independence between outcomes and hidden states, 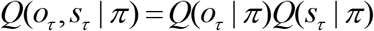, the resulting belief update equations would correspond exactly to a variational message passing algorithm (Dauwels 2007) applied to a model with missing data.

When the expectation is taken with respect to the approximate posteriors, the marginalisation implicit in this definition ensures that

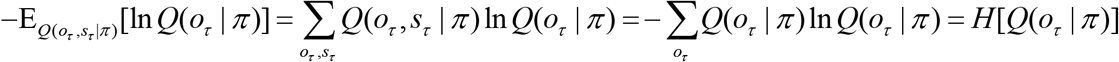

If we write out the generative model in full, and substitute this (omitting constants) into Equation 3, we can use the same implicit marginalisation to write:

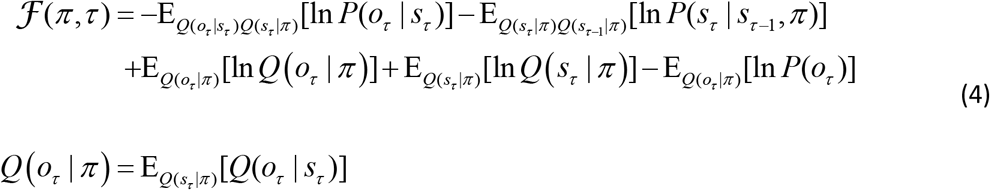

The implicit generative model now incorporates a prior over observations. This means that the generative model is replaced with that shown on the right of Figure 1:

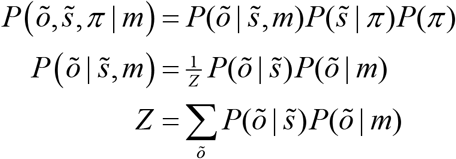

Here, we have defined the distribution over states and observations in terms of two independent factors, a likelihood, and a prior over observations; i.e. preferred observations conditioned on the model. For simplicity, we will omit the explicit conditioning on *m*, so that 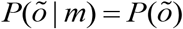. For past states, this distribution is flat. Crucially, this means the generalised free energy reduces to the variational free energy for outcomes that had been observed in the past. Separating out contributions from the past and the future, we are left with the following:

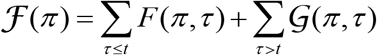

Unlike *G* (the expected free energy), 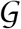 is the free energy of the expected future. We can rearrange Equation 4 (for future states) in several ways that offer some intuition for the properties of the generalised free energy.

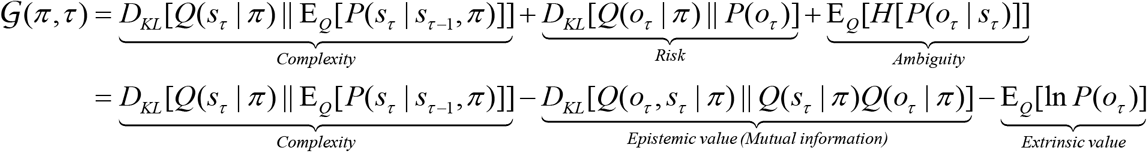

To obtain the mutual information term, we have used the relationship 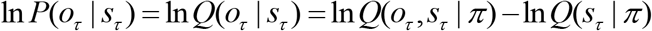. The imperative to maximise the mutual information (Barlow 1961; Barlow 1974; Linsker 1990; Optican and Richmond 1987) can be interpreted as an epistemic drive (Denzler and Brown 2002). This is because policies that (are believed to) result in observations that are highly informative about the hidden states are associated with a lower generalised free energy. As a KL-Divergence is always greater than or equal to zero, the second equality indicates that the free energy of the expected future is an upper bound on expected surprise.

To find the belief update equations for the policies, we take the variational derivative of the generalised free energy with respect to the posterior over policies, and set the result to zero in the usual way:

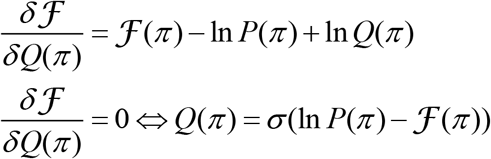

At time *τ*=0, no observations have been made, and the distribution above becomes a prior. When this is the case, 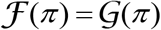, so the prior over policies is:

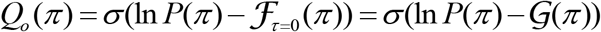

If we take the variational derivative of Equation 4 with respect to the hidden states:

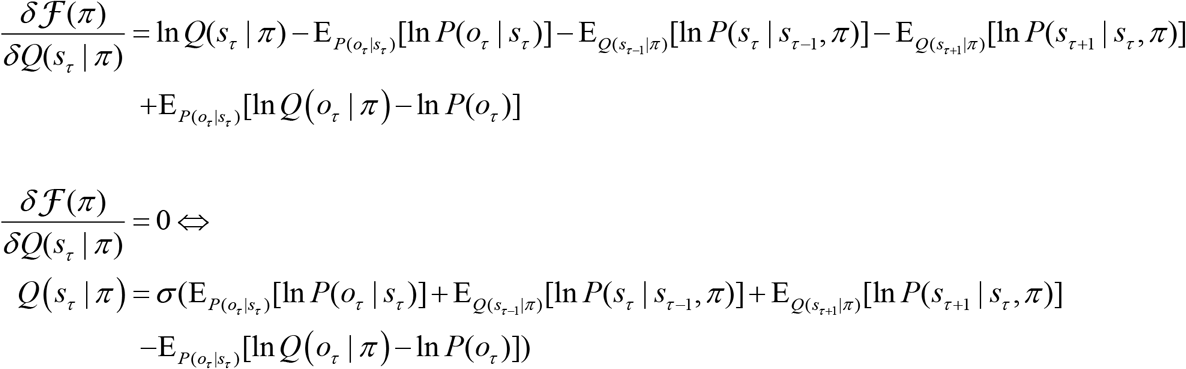

The derivative of 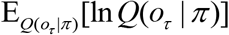 is a little complicated, so this is presented step by step in Appendix B. The hidden state update has a different interpretation in the past compared to the future:

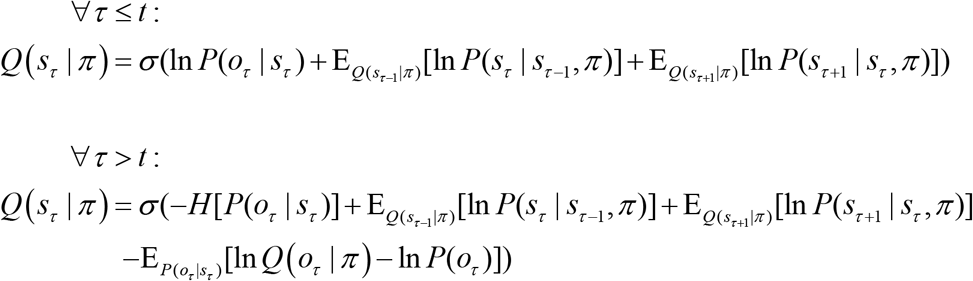

The final term for future beliefs implies that future states are considered more probable if they are expected to be similar to those that generate preferred outcomes. In other words, there is an optimistic distortion of beliefs about the trajectory into the future.

### Summary

We have introduced a generalised free energy functional that is expressed as a functional of beliefs about data. The variational free energy can be seen as a special case of this generalised functional, when beliefs about outcomes collapse to delta functions. When we derive update equations (Figure 3, pink panels) under this functional, the updates look very similar to those based on the variational free energy approach. An important difference between the two approaches is that we have now included the prior probability of outcomes in the generative model. This has no influence over beliefs about the past, but distorts beliefs about the future in an optimistic fashion. This formulation generalises not only the standard active inference formalism, but also active data selection or sensing approaches in machine learning (MacKay 1992) and computational neuroscience (Yang et al. 2016b). See Appendix A for a discussion of the relationship between these.

## 4 Comparison of active inference under expected and generalised free energy

The generalised free energy has the appeal that belief updating and policy selection both minimise the same objective function. The temporal symmetry of this free energy ensures that it is a path integral through time. The use of this integral to evaluate the probability of a set of plausible trajectories resembles the use of Lagrangians in physics, under Hamilton’s principle of stationary action. In contrast, formulations of active inference to date have required two different quantities (the variational free energy and the expected free energy respectively) to derive these processes. Although the form of belief updating is the same, the belief updates resulting from the use of a generalised free energy are different in subtle ways. In this section, we will explore these differences, and show how generalised active inference reproduces the behaviours illustrated in our earlier papers.

The notable differences between the updates are found in the policy prior, the treatment of outcomes, and the future hidden state updates. The prior over policies is very similar in both formulations. The expected and generalised free energy (at *τ* = 0) differ only in that there is an additional complexity term in the latter. This has a negligible influence on behaviour, as the first action is performed *after* observations have been made at the first time step. At this point, the posterior belief about policies is identical; as the variational free energy supplies the missing complexity term. Although the priors are different, both in form and motivation, the posterior beliefs turn out to be computed identically. Any difference in these can be attributed to the quantities used to calculate them; namely, the outcomes and the hidden states.

Outcomes in the generalised formulation are represented explicitly as beliefs. This means that the prior over outcomes is incorporated explicitly in the generative model. There are two important consequences of this. The first is that the posterior beliefs about outcomes can be derived in a parsimonious way, without the need to define additional prior distributions. The second is that hidden state beliefs in the future are biased towards these preferred outcomes. A prior belief about an outcome at a particular time point thus distorts the trajectory of hidden states at each time point reaching back to the present. In addition to this, beliefs about hidden states in the future acquire an ‘ambiguity’ term. This means that states associated with an imprecise mapping to sensory outcomes are believed less likely be inferred. In summary, not only are belief trajectories drawn in optimistic directions, they also tend towards states that offer informative observations.

To make the abstract considerations above a little more concrete, we have employed an established generative model that has previously been used to demonstrate epistemic behaviours under active inference (Friston et al. 2015). This is a T-maze task (Figure 4), in which an agent decides between (temporally deep) policies. In one arm, there is an unconditioned (rewarding) stimulus. In another, there is no stimulus, and this condition is considered aversive. In the final arm, there is always an instructional or conditioned stimulus that indicates the arm that contains the reward. The starting location and the location of the conditioned stimulus are neither aversive nor rewarding.

**Figure 4 -.**
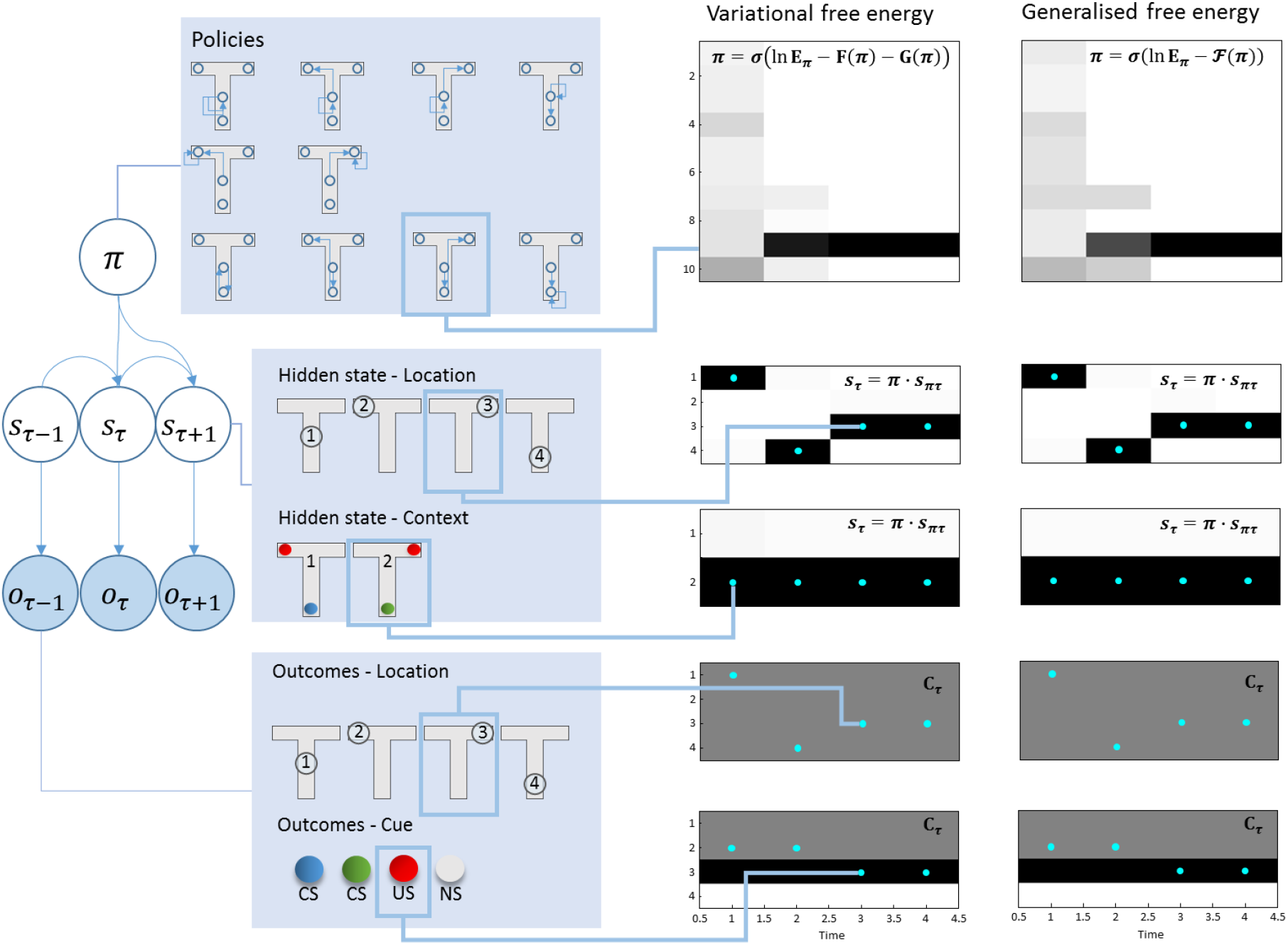
T-maze simulation. The *left* part of this figure shows the structure of the generative model used to illustrate the behavioural consequences of each set of update equations. We have previously used this generative model to address exploration and exploitation in two-step tasks; further details of which can be found in Friston et al. (2015). In brief, an agent can find itself in one of four different locations, and can move among these locations. Locations 2 and 3 are absorbing states, so the agent is not able to leave these locations once the have been visited. The initial location is always 1. Policies define the possible sequences of movements the agent can take throughout the trial. For all 10 available policies, after the second action, the agent stays where it is. There are two possible contexts: the unconditioned stimulus (US) may be in the left or right arm of the maze. The context and location together give rise to observable outcomes. The first of these is the location, which is obtained through an identity mapping from the hidden state representing location. The second outcome is the cue that is observed. In location 1, a conditioned stimulus (CS) is observed, but there is a 50% chance of observing blue or green, regardless of the context, so this is uninformative (and ambiguous). Location 4 deterministically generates a CS based on the context, so visiting this location resolves uncertainty about the location of the US. The US observation is probabilistically dependent on the context. It is observed with a 90% chance in the left arm in context 1, and a 90% chance in the right arm in context 2. The *right* part of this figure compares an agent that minimises its variational free energy (under the prior belief that it will select policies with a low expected free energy) with an agent that minimises its generalised free energy. The upper plots show the posterior beliefs about policies, where darker shades indicate more probable policies. Below these, the posterior beliefs about states (location and context) are shown, with blue dots superimposed to show the true states used to generate the data. The lower plots show the prior beliefs about outcomes (i.e., preferences), and the true outcomes (blue dots) the agent encountered. Note that a US is preferred to either CS, both of which are preferable to no stimulus (NS).

As Figure 4 shows, regardless of the active inference scheme we use, the agent first samples the unrewarding, but epistemically valuable, uncertainty resolving cue location. Having resolved uncertainty about the context of the maze, the agent proceeds to maximise its extrinsic reward by moving to the reward location. Although the most striking feature of these simulation results is their similarity, there are some interesting differences worth considering. These are primarily revealed by the beliefs about hidden states over time. Under each of the schemes presented here, there exist a set of (neuronal) units that encode beliefs about each possible state. For each state, there are units representing the configuration of that state in the past and future, in addition to the present. The activity in these units is shown in Figure 5. The differences here are more dramatic than in the subsequent behaviours illustrated in Figure 4. At the first time step (column 1), both agents infer that they will visit location 4 at the next time, resolving uncertainty about the context of the maze. From this future point onwards, however, the beliefs diverge. This can be seen clearly in the lower rows of column 1; the beliefs about the future at the first time step. The agent who employs expected free energy believes they will stay in the uncertainty resolving arm of the maze, while the generalised agent believes they will end up in one of the (potentially) rewarding arms. Despite a shared proximal belief trajectory, the distal elements of the two agents’ paths are pulled in opposite directions. As each future time point approaches, the beliefs about that time begin to converge – as observations become available.

**Figure 5 -.**
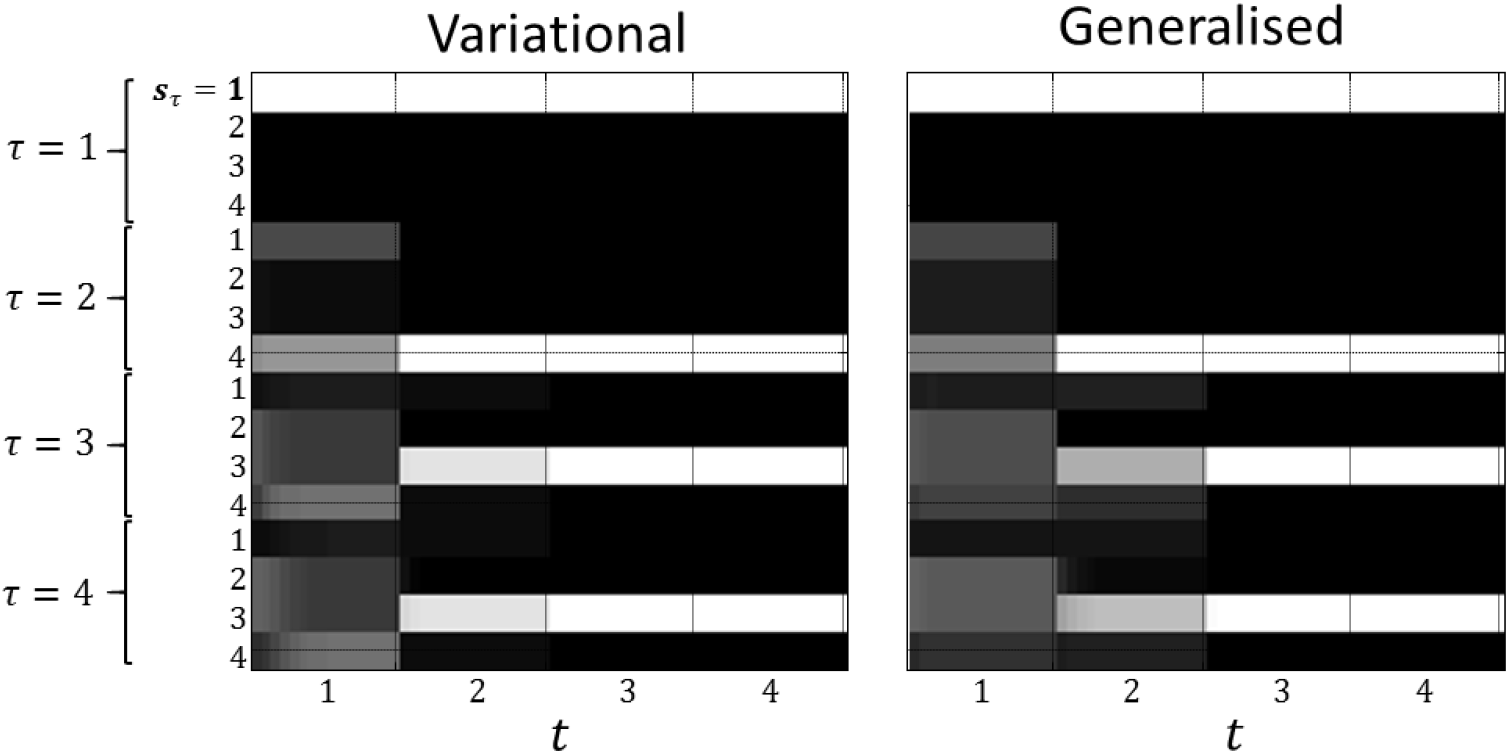
Optimistic distortions of future beliefs. These raster plots represent the (Bayesian model average of the) approximate posterior beliefs about states (specifically, those pertaining to location). At each time step *t*, there is a set of units encoding beliefs about every other time step *τ* in the past and future. The evolution of these beliefs is reflected the evidence a cumulation or belief updating of approximate posterior expectations, with lighter shades indicating more probable states

## 5. Conclusion

The generalised free energy introduced in this paper provides a new perspective on active inference. It unifies the imperatives to minimise variational free energy with respect to data, and expected free energy through model selection, under a single objective function. Like the expected free energy, this generalised free energy can be decomposed in several ways; giving rise to familiar information theoretic measures and objective functions in Bayesian reinforcement learning. Generalised free energy minimisation replicates the epistemic and reward seeking behaviours induced in earlier active inference schemes, but prior preferences now induce an optimistic distortion of belief trajectories into the future. This allows beliefs about outcomes in the distal future to influence beliefs about states in the proximal future and present. That these beliefs then drive policy selection suggests that, under the generalised free energy formulation, the future can indeed cause the past.

## Appendix A Active data selection

Active data selection has been a topic of interest in both neuroscience and machine learning for a number of years (Krause 2008). Several different approaches have been taken to define the best data to sample (Settles 2010), and the optimal experiments to perform to do this (Daunizeau et al. 2011). This appendix addresses the relationship between the future components of the expected free energy and established methods. Writing in full, the (negative) free energy of the expected future is

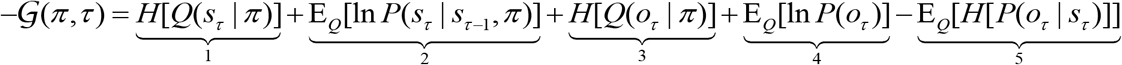

Under active inference, the above functional is maximised. If we were to use only term 3, this maximisation reduces to ‘uncertainty sampling’ (Hwa 2004; Lewis and Gale 1994; Shewry and Wynn 1987). This involves (as the name suggests) selecting the data points about which uncertainty is highest. A problem with this approach is that it may favour the sampling of ambiguous (uninformative) data. A more sophisticated objective function includes both 3 and 5 (Denzler and Brown 2002; Lindley 1956; MacKay 1992; Yang et al. 2016a). This means that uncertain data points are more likely to be sampled, but only if there is an unambiguous mapping between the latent variable of interest and the data. Term 4 is a homologue of expected utility (reward) in reinforcement learning (Sutton and Barto 1998), and is an important quantity in sequential statistical decision theory (El-Gamal 1991; Wald 1947). Terms 1 and 2 together contribute to an ‘Occam factor’ (Rasmussen and Ghahramani 2001); a component of some previously used objective functions (MacKay 1992).

All of these quantities are emergent properties of a system that minimises its expected free energy. In the schemes mentioned above, the quantities were pragmatically selected to sample data efficiently. Here, they can be seen as special cases of the free energy functional used to define the active inference or sensing that underwrites perception (Friston et al. 2012b; Gregory 1980).

## Appendix B Variational derivative of expected marginal

Below are the steps taken to obtain the variational derivative of an expected marginal. This is needed for the hidden state update equations under the generalised free energy.

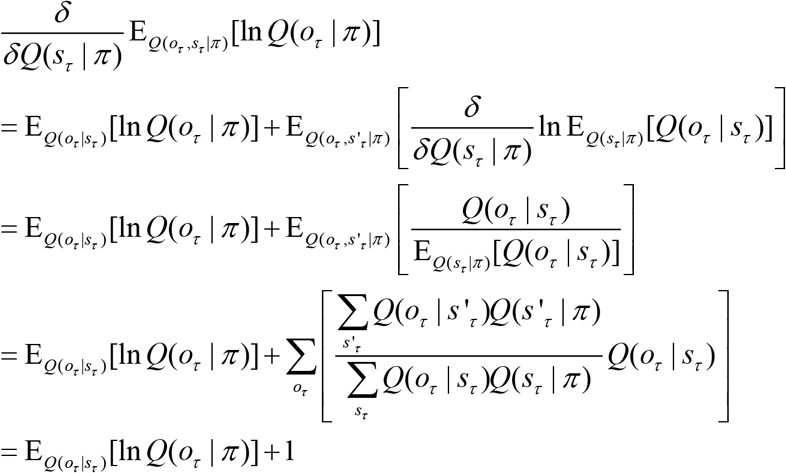

In the update equations, we can omit the constant 1.

## Acknowledgements

TP is supported by the Rosetrees Trust (Award Number 173346). KJF is a Wellcome Principal Research Fellow (Ref: 088130/Z/09/Z). The authors thank Dimitrije Markovic for his insightful comments on the manuscript.

